# An intact oculomotor neural circuit in congenital blindness

**DOI:** 10.1101/2023.03.24.533989

**Authors:** Cemal Koba, Alessandro Crimi, Olivier Collignon, Emiliano Ricciardi, Uri Hasson

## Abstract

In the past three decades, multiple studies revealed that congenital blindness is associated with functional and structural reorganization in early visual areas and its interaction with other neural systems. Among the most reproducible findings is the weaker connectivity between the visual and sensorimotor cortices, which in sighted individuals plays a role in eye-motor coordination. Here we demonstrate an important exception to this reorganization phenomena: we find that in congenitally blind individuals (as for normally sighted ones), spontaneous, non-controlled eye movements correlate with connectivity between visual and sensorimotor cortices. Furthermore, using time-lagged regression, we show that eye movements drive activity in the visual cortex, which subsequently drives sensorimotor activity. Remarkably, this phenomenon persists even though blind participants often exhibit unsynchronized eye movements and cannot sense or regulate their eye positions. These findings provide evidence of a preserved, “non-functional” connectivity in congenitally blind individuals and reveal a firm, hard-wired constraint on connectivity that remains immune to any reorganization phenomena produced by lack of visual experience and oculomotor control.

## Introduction

Congenital blindness (CB) inevitably causes profound changes in the control of eye movements. Previous studies documented slow, irregular oscillations that are sometimes uncoordinated between the two eyes (e.g., Kompf & Piper, 1987; Leigh & Zee, 1980). These studies also showed that CB individuals are typically unaware of their spontaneous eye movements, unable to sense eye position, and cannot voluntarily initiate saccades toward a specific direction. In contrast to sighted groups, people with CB manifest slow eye drifts after rapid eye movements or saccades and lack a normal vestibulo-ocular reflex (in which the eyes move in a direction opposite to head movement). This suggests the absence of afferent information from the vestibular to the oculomotor systems (Kompf & Piper, 1987). As such, control and feedback loops related to the oculomotor system are strongly altered when visual input is absent since birth.

In sighted groups, endogenously-driven oculomotor patterns impact the topography and topology of functional brain networks. Specifically, our group previously characterized the neural correlates associated with spontaneous, non-directed eye movements by extracting EPI signal from the eye orbit (EO-EPI) area (Koba et al., 2021). This EO-EPI signal mirrors simultaneously collected eye-tracking data and was here used as a seed time series in modeling whole-brain resting state (RS) data. The analyses showed that in sighted individuals, spontaneous eye movements are specifically associated with bilateral activity in sensorimotor regions (pre- and post-central gyri and central sulcus, including the frontal eye fields), supplementary motor area, and cerebellum. Consistent with this, partialling out the variance related to eye movements from the RS data reduced connectivity between visual cortex (VisCtx) and sensorimotor cortex (SMCtx), thus confirming that oculomotor-related contributions form an important component of resting state network topology. The fact that eye movements are related to VisCtx-SMCtx connectivity is per se non-surprising, since these two areas support visuo-motor processing tasks (e.g., Bernardi et al., 2013).

Given both the lack of a lifelong visual input and a physiological oculomotor control since birth, the characterization of whether the altered spontaneous eye movement patterns in CB may affect functional brain networks could provide a deeper understanding of the principles that determine organized patterns of RS connectivity. In particular, we asked whether altered eye movements in CB would still consistently impact the functional connectivity between the sensorimotor system and visual areas that are linked to oculomotor activity. Consistency would be indicated by overlapping patterns at the group level, reflecting an eye movement-dependent, but experience-independent role in mediating resting-state brain connectivity. In this case, the altered spontaneous eye movements in CB would still reliably affect sensorimotor-to-visual connectivity, thus indicating an intrinsic constraint on connectivity that remains immune to any reorganization phenomena due to the lack of visual experience and ensuing oculomotor control. A plausible alternative hypothesis is that eye movement-related networks will associate with different connectivity patterns across CB individuals. In this case, the disorganized eye movements in CB would heterogeneously impact sensorimotor-to-visual connectivity depending on individual (e.g., blindness etiology) and experience-dependent factors.

Prior research presents mixed findings in relation to this question. The observation that eye movements correlate with activity in VisCtx and SMCtx already in fetuses suggests that visual experience is unnecessary for this network to develop (Schöpf et al., 2014). Consistent with this finding, Sen et al. (2022) quantified inter-subject variability associated with functional connectivity (FC) among different brain areas in both sighted and CB and concluded that there are limited neoplastic changes in the VisCtx-to-SMCtx connectivity of CB individuals. Notably, while the strength profile of resting-state connectivity was often more heterogeneous in CB than Sighted, connectivity between VisCtx and SMCtx showed less variability for CB. Thus, although a broader functional reorganization process at a whole brain level occurs due to the congenital loss of visual input (e.g., Bock & Fine, 2014; Castaldi et al., 2020; Voss, 2019), the sensorimotor system related to oculomotor activity appears to develop early in life and to be less affected by (the lack of) visual experience-related changes. However, contrary to those findings, it has been reliably demonstrated that CB individuals show reduced RS-FC between visual and sensorimotor systems, suggesting alterations in the visuomotor circuit related to eye movement. This is one of the most notable differences in connectivity already identified by early studies (Liu et al., 2007) and reviews (Bock & Fine, 2014).

In conclusion, finding that eye movements synchronize visual and sensorimotor cortices in blindness since birth would provide evidence for an intact circuit that serves no recognizable function or computation, and may - in this sense - reflect the presence of “nonfunctional connectivity” in brain networks. In contrast, if eye movements do not synchronize FC between these regions in CB, this would suggest this specific circuit undergoes neuroplastic changes due to lack of functional purpose, and would present an alternative explanation for the weaker connectivity between VisCtx and SMCtx repeatedly documented in prior work.

## Methods

### Dataset

Complete details of the dataset and imaging parameters are given in Pelland et al. (2017), and here we report only the main details. The entire dataset includes 50 participants that participated in a single five-minute functional MRI run (136 volumes). Participants were instructed to keep their eyes closed, relax, and not think about anything in particular. Functional time series were acquired using a 3-T TRIO TIM (Siemens) equipped with a 12-channel head coil. Multislice T2*-weighted fMRI images were obtained with a gradient echo-planar sequence using axial slice orientation; repetition time (TR) 2,200 ms; echo time (TE) 30 ms; functional anisotropy (FA) 90°; 35 transverse slices; 3.2 mm slice thickness; 0.8 mm gap; field of view (FoV) 192 × 192 *mm*^2^; matrix size 64 × 64 × 35; voxel size 3 × 3 × 3.2 *mm*^3^. A structural T1-weighted 3D magnetization prepared rapid gradient echo sequence (voxel size 1 × 1 × 1.2 mm^3^; matrix size 240 × 256; TR 2,300 ms; TE 2.91 ms; TI 900 ms; FoV 256; 160 slices) was also acquired for all participants. All of the procedures were approved by the Research Ethic and Scientific Boards of the Centre for Interdisciplinary Research in Rehabilitation of Greater Montreal and the Quebec Bio-Imaging Network. Experiments were undertaken with the understanding and written consent of each subject.

The 50 participants included 14 CB individuals (9 males and 5 females, with a mean age of 44.93 ± 11.19 years and 12 right-handed and 2 ambidextrous), as well as sighted controls for the congenitally blind participants, late-blind participants and sighted controls for the late blind. In this study, we analyze data for the CB only, since the Sighted participants were blindfolded during the scan and were not used as a comparison group due to the differences in eye movements between eyes-open and eyes-closed states in Sighted individuals (Pelland et al., 2017). In a few cases, when required, we used as a reference previously analyzed data from a resting-state study where sighted controls fixated on a screen center with open eyes (see details below).

### Data Preprocessing

The fMRI data were made available using the Brain Imaging Data Structure (BIDS, Gorgolewski et al., 2016) format. Along with the folder and naming standardization, the first four volumes of functional runs were removed to avoid stabilization artifacts (leaving 132 volumes in total), and the resolution of the functional runs was interpolated to 3 × 3 × 4 mm^3^. The providers of the data confirmed that no other preprocessing steps were applied during the standardization procedure.

The results reported in this manuscript are based on initial preprocessing that we performed using *fMRIPrep* 20.2.1 (Esteban, Markiewicz, et al., 2018; Esteban, Blair, et al., 2018; RRID:SCR_016216), which is based on *Nipype* 1.5.1 (Gorgolewski et al., 2011; Gorgolewski et al., 2018; RRID:SCR_002502). The preprocessing steps applied by *fMRIPrep* are listed below:

#### Anatomical data preprocessing

A total of 1 T1-weighted (T1w) image were found within the input BIDS dataset. The T1-weighted (T1w) image was corrected for intensity non-uniformity (INU) with `N4BiasFieldCorrection` (Tustison et al., 2010), distributed with ANTs 2.3.3 (Avants et al., 2008) (RRID:SCR_004757), and used as T1w-reference throughout the workflow. The T1w-reference was then skull-stripped with a *Nipype* implementation of the `antsBrainExtraction.sh` workflow (from ANTs), using OASIS30ANTs as target template. Brain tissue segmentation of cerebrospinal fluid (CSF), white-matter (WM) and gray-matter (GM) was performed on the brain-extracted T1w using `fast` (Zhang et al., 2001) (FSL 5.0.9, RRID:SCR_002823). Volumebased spatial normalization to one standard space (MNI152NLin2009cAsym) was performed through nonlinear registration with `antsRegistration` (ANTs 2.3.3), using brain-extracted versions of both T1w reference and the T1w template. The following template was selected for spatial normalization: *ICBM 152 Nonlinear Asymmetrical template version 2009c* (Fonov et al., 2009, RRID:SCR_008796; TemplateFlow ID: MNI152NLin2009cAsym).

#### Functional data preprocessing

The following pre-processing was performed on the RS data of each subject. First, a reference volume and its skull-stripped version were generated using a custom methodology of *fMRIPrep*. Susceptibility distortion correction (SDC) was omitted. The Blood-oxygen-level-dependent (BOLD) reference volume was then co-registered to the T1w reference using `flirt` (Jenkinson & Smith, 2001) with the boundary-based registration (Greve & Fischl, 2009) cost-function. Co-registration was configured with nine degrees of freedom to account for distortions remaining in the BOLD reference. Head-motion parameters with respect to the BOLD reference (transformation matrices, and six corresponding rotation and translation parameters) were estimated before any spatiotemporal filtering, using `mcflirt` (Jenkinson et al., 2002)(FSL 5.0.9). The BOLD times-series were resampled onto their original, native space by applying the transforms to correct head motion. These resampled BOLD time series will be called *preprocessed BOLD in the original space*, or just *preprocessed BOLD*. The BOLD time series were then resampled into standard space, generating a *preprocessed BOLD run in MNI152NLin2009cAsym space*. All resamplings can be performed with *a single interpolation step* by composing all the pertinent transformations (i.e. head-motion transform matrices, susceptibility distortion correction when available, and co-registrations to anatomical and output spaces). Gridded (volumetric) resamplings were performed using `antsApplyTransforms` (ANTs), configured with Lanczos interpolation to minimize the smoothing effects of other kernels (Lanczos, 1964).

Several confounding time series were calculated based on the *preprocessed BOLD:* framewise displacement (FD), DVARS and three region-wise global signals. FD was computed using two formulations following Power (absolute sum of relative motions, Power et al., 2014) and Jenkinson (relative root mean square displacement between affines, Jenkinson et al., 2002). FD and DVARS are calculated for each functional run, both using their implementations in *Nipype* following the definitions by Power et al. (2014). The three global signals are extracted within the CSF, the WM, and the whole-brain masks. The head-motion estimates calculated in the correction step were also placed within the corresponding confounds file. The confound time series derived from head motion estimates and global signals were expanded with the inclusion of temporal derivatives and quadratic terms for each (Satterthwaite et al., 2013). Frames that exceeded a threshold of 0.5 *mm* FD or 1.5 standardized DVARS were annotated as motion outliers.

Many internal operations of *fMRIPrep* use *Nilearn* 0.6.2, RRID:SCR_001362 (Abraham et al., 2014), mostly within the functional processing workflow. For more pipeline details, see [the section corresponding to workflows in *fMRIPrep*’s documentation](https://fmriprep.readthedocs.io/en/latest/workflows.html).

Copyright Waiver: The above boilerplate text was automatically generated by fMRIPrep with the express intention that users should copy and paste this text into their manuscripts *unchanged*. It is released under the [CC0](https://creativecommons.org/publicdomain/zero/1.0/) license.

After the above procedure applied by *fMRIprep*, we applied band-pass filtering (0.01–0.1 Hz) and cleaned the data from 6 main motion parameters, mean WM and CSF signals, and FD using *AFNI’s* `3dDeconvolve` (Cox, 1996). The resulting dataset was smoothed with 6mm full-width half maximum (FWHM) kernel with the `3dBlurToFWHM` function of the same software. These smoothed residuals were considered as the resting-state data to be used in all subsequent analyses.

All the figures in this paper were generated using *Nilearn* (Abraham et al., 2014), *MATLAB version 9.10.0.1613233 (R2021a)*, 2021, or *BrainNet Viewer* (Xia et al., 2013).

#### Creating the Eye Orbit EPI (EO-EPI) regressors

The Eye-Orbit (EO) area was marked using *MRICRON* (Rorden et al., 2007) on the common anatomical template *MNI152 NLin 2009c Asym* (Supplementary Figure 1). The EO region of interest (ROI) covered the entire eye orbit. The mask was drawn on the MNI template and then applied to all subjects regardless of their eye vitreous size or availability. The resulting binary ROI mask was resampled to the resolution of the functional runs and no further processing was needed because all the resting-state data were already aligned to this common space. The resampled mask was used to extract the mean resting-state signal from the eye regions (denoted as EYE_*raw*_) for each subject. EO-EPI time series extracted from both eyes should be strongly correlated. As reported in the Results, this was the case for the sighted controls who participated in the current study with their eyes blindfolded. While the correlation was low or negative for some CB participants, we still decided to average the time series in such cases. Our motivation was that averaging emphasizes epochs of coordinated movement. However, we also considered that neuroplastic changes in the blind may produce divergent ocular-cortical modulation for each eye, as we document for two cases in the Supplementary Materials. The average time series were also convolved with a basis HRF function using *AFNI’s* `waver` command, producing EYE_*conv*_. In separate analyses, we used either EYE_*raw*_ or EYE_*conv*_ as “seed” regressors to identify brain areas correlated with the EO-EPI signal.

### Statistical Inference of fMRI Analyses

#### Correlates of EO-EPI regressors

We conducted a whole-brain single-voxel regression for each participant using a simple univariate linear model. In this model, EYE_*raw*_ was the independent variable and voxel-wise resting-state data was the dependent variable (note that nuisance factors were removed during preprocessing). The significance of the beta coefficients was determined at the group level using *FSL’s* `randomize` function (Winkler et al., 2014). This function applies a one-sample T-test and determines the significance threshold through 10,000 permutations and threshold-free cluster enhancement. The age and sex of each subject were accounted for in the group-level statistics by inserting them as regressors of no interest. The same procedure was repeated using EYE_*conv*_ as the seed time series.

#### The impact of eye movement on connectivity

To create functional connectivity networks, we used a resting-state functional connectivity parcellation based on 400 ROIs grouped by 7 networks: Visual, Somatomotor, Dorsal Attention, Ventral Attention, Limbic, Frontoparietal, and Default Mode (Glen et al., 2021; Schaefer et al., 2018). This parcellation was also aligned to the template we used (*MNI152NLin2009cAsym*). We extracted the mean times-series from each ROI, for the two types of spatially smoothed resting-state data we derived (one typical, and the other with EO-EPI *EYE_raw_* regressed out).

To determine the effect of partialling out *EYE_raw_* from the resting-state data on functional connectivity networks, as a first step we constructed two variants of the 400-by-400 connectivity matrix. Each was created by correlating the mean resting-state time series extracted from the 400 ROIs using the Schaefer et al. (2018) atlas. One matrix was based on the original resting-state data, and the other used the resting-state data that was cleaned of EYE_*raw*_. Ultimately this produced two correlation matrices for each participant.

To identify if there were clusters of regions whose connectivity changed after the removal of the EO-EPI data, the two 400 × 400 connectivity matrices were compared using a difference-network analysis using the Network-Based Statistics Toolbox (NBS, Zalesky et al.,2010), implemented in MATLAB (*MATLAB version 9.10.0.1613233 (R2021a)*, 2021). This analysis quantifies the likelihood of finding a cluster of nodes all of which are more weakly connected in one condition than the other. Initial thresholding (a hyper-parameter) ensures that only strong connections are considered. We used primary t-thresholds of 7.5 and of 5.0, 10,000 permutations, and an alpha value of 0.05.

#### Effective connectivity

To further examine those connections that were weaker in resting-state networks with removed EO-EPI (specifically, the connection between the right visual cortex and sensorimotor area) and the EYE_*raw*_, we used an effective connectivity model based on Granger Causality. The goal was to understand the causal interaction between eye movements, cortical signals, and the direction of the connection between the occipital and sensorimotor cortices. The analysis was applied via a custom script written in Python (Van Rossum & Drake, 2009), using the *statsmodels* library (Seabold & Perktold, 2010). Before performing the analysis for effective connectivity, a preliminary step was applied to ensure the stationarity of the time series. More specifically, the augmented Dickey-Fuller test was employed (Dickey & Fuller, 1979). Once the stationarity of the signal was confirmed, we implemented pairwise Granger causality models with an inclusive range of time lag from 1 to 5. The multivariate approach was not used given the small number of regions of interest. After estimating the autoregressive model, F-statistics were computed and converted into p-values. For each subject, each time lag (from 1 to 5 TR) and each causality test (visual-sensorimotor, visual-EYE_*raw*_; sensorimotor-EYE_*raw*_, sensorimotor-visual; EYE_*raw*_-visual, EYE_*raw*_-sensorimotor), p-values were calculated. To avoid sensitivity to outliers, the median across the entire dataset was computed and reported (following Duggento et al., 2018).

## Results

### Temporal characteristics of EO-EPI time series in blind

As expected, CB presented ocular movements with different characteristics from those of Sighted. In the current dataset, this was evident in several ways. First, we computed the frequency characteristics of the EO-EPI EYE_*raw*_ time series in CB and compared those to data from Sighted that we used in our previous study (Koba et al., 2021), based on the SleepyBrain dataset (Nilsonne et al., 2016). Example time series from CB and Sighted are presented in Figure 1A. Visual inspection suggests that in CB, the spectral power is more spread over the frequency range. There was also higher power for CB in the lower (<0.01 H*z*) and higher (0.05-0.1 H*z*) frequencies (Figure 1C). EO-EPI data of CB also tended to have a larger variance per time series (mean = 0.04 ± 0.01), as compared to that of Sighted (mean = 0.03 ±0.001, *t* = 3.79, p<0.001, Figure 1B). As shown in Supplementary Materials, EO-EPI time series from each eye were positively correlated in Sighted but less in CB (Supplementary Figure 3). In some cases, CB participants had one eye with more pronounced dynamics than the other (Supplementary Figure 4).

**Figure 1.**
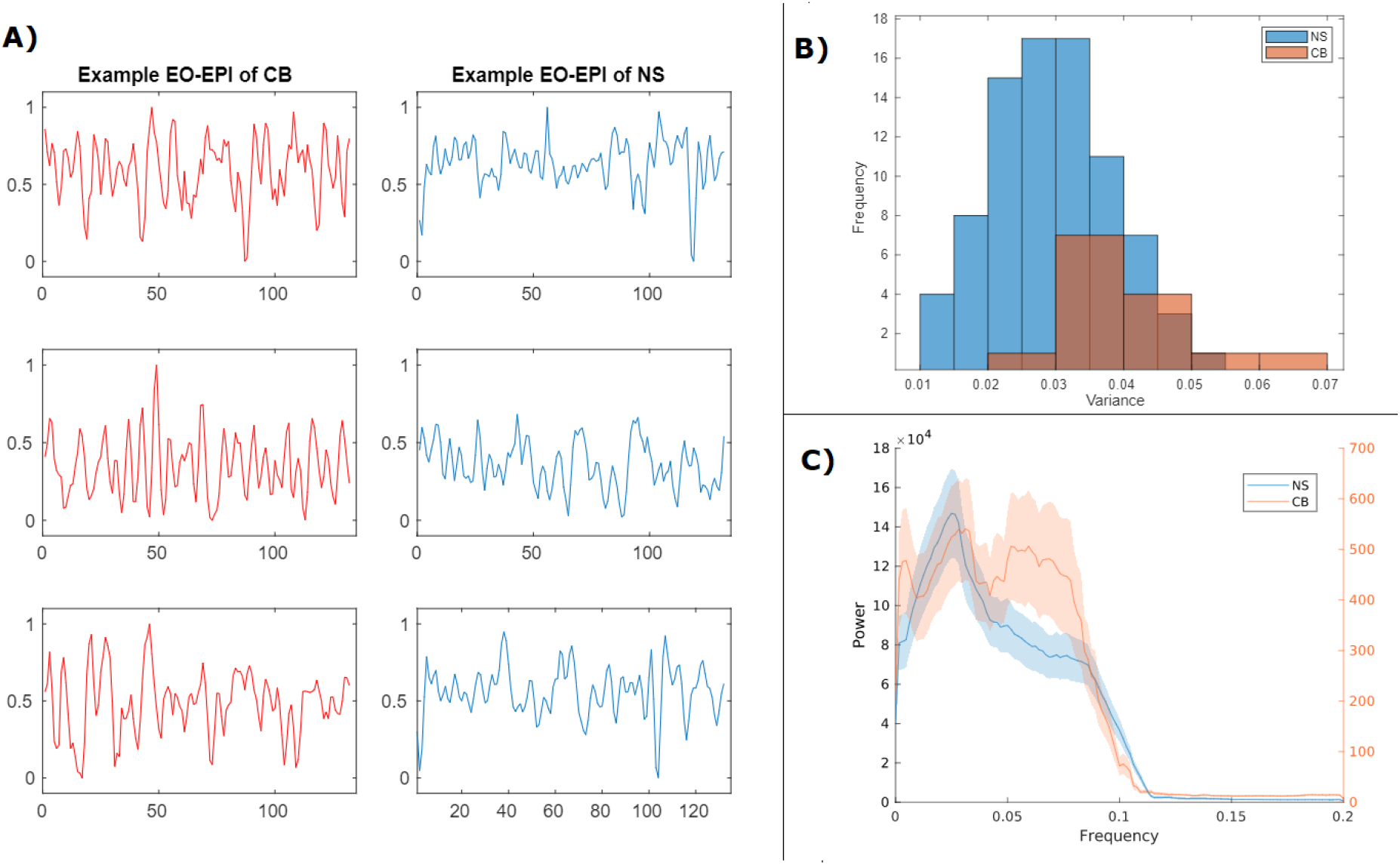
Comparison of EO-EPI time series in CB and Normally Sighted (NS). Series from both groups were normalized into the range of 0 to 1. **A)** Example EO-EPI time series from CB (left column) and Sighted (right column). **B)** Histogram of the variances of EO-EPI time series. **C)** Power-frequency distribution of EO-EPI time series (in H*z*).

### Correlates of EO-EPI regressor

Correlates of EYE_*conv*_ and EYE_*raw*_ on the whole-brain level were determined via a one-sample T-test whose sampling distribution was determined using 10,000 permutations, and the results were corrected with FWE of *α* = 0.05 using TFCE. Brain regions positively related to EYE_*raw*_ are shown in Figure 2. While a positive relationship is found with bilateral superior frontal sulci, posterior cingulate, lateral occipital cortices, and cerebellum, a negative relationship is visible with the sensorimotor area. Correlates of EYE_*conv*_ captured a different, more limited pattern (Supplementary Figure 2), showing a positive relationship in the head of the right caudate nucleus and a negative relationship for the thalamus and visual cortex. A complete listing of these regions according to the Anatomical Automatic Labeling 2 template (AAL2, Rolls et al., 2015), Desikan-Killiany (Desikan et al., 2006), and Harvard-Oxford (RRID:SCR_001476) atlases are reported in Supplementary Table 1 and 2.

**Figure 2.**
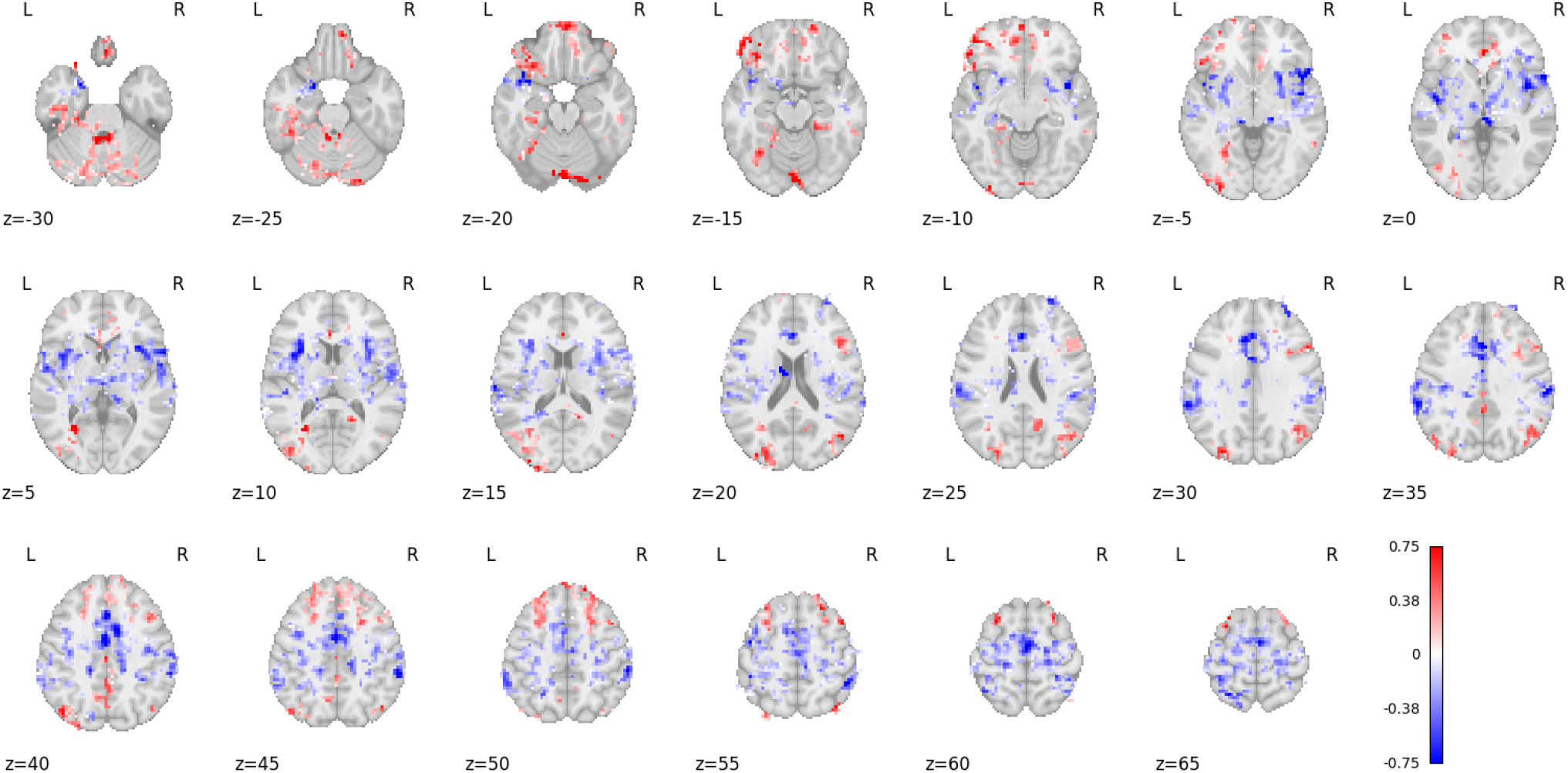
Brain areas where BOLD activity correlated with eye movements in congenitally blind. The figure shows correlates of EYE_*raw*_, p<0.05, TFCE-corrected for multiple comparisons. Warm colors represent positive correlations; cold colors represent negative ones.

We compared the spatial distribution of results to that found for Sighted individuals in our previous study (Koba et al., 2021), which included 83 participants. The spatial distribution of activity identified for EYE_*raw*_ showed higher spatial overlap than that observed for EYE_*conv*_, as evidenced by Dice coefficients of 0.25 and 0.09, respectively. For this reason, we proceed with the EYE_*raw*_ in subsequent analyses. Supplementary Figure 5 presents the spatial overlap between the statistical maps produced for CB and Sighted.

As indicated above, while we averaged the two EO-EPI time series per participant to identify time points of coordinated eye movements, in some CB participants, the correlation was low. For two CB participants, we conducted an exploratory analysis, where connectivity maps were computed separately for each eye. The results are presented in Supplementary Figure 6. As shown in the figure, in the cases where the correlation was low, each eye appeared to pick up different brain networks.

#### Functional connectivity

To evaluate the impact of EO-EPI on functional connectivity networks, we created two correlation matrices for each participant. The matrices, each with a dimension of 400 × 400 corresponding to the number of ROIs in the parcellation scheme adopted, were derived from the regular RS data and EO-EPI-removed RS data, respectively.

To evaluate larger-scale topographical differences in connectivity, we used the Network-Based Statistics (NBS) toolbox to identify “difference networks” consisting of a continuous set of interconnected edges with lower values in one condition than another. For the EO-EPI-removed networks, we found a significantly reduced connectivity between the VisCtx and SMCtx (Supplementary Figure 7). The MNI coordinates of these nodes are *24, −99, 7* for the VisCtx (corresponding to BA17/18 according to the Juelich atlas, Eickhoff et al., 2007) and *22, −35, 71* for the SMCtx (corresponding to BA3b and BA4a according to the Juelich atlas). To determine the absolute correlation values, we extracted time series from each of these ROIs and computed the Z-transformed Pearson correlations. Mean Z-transformed connectivity values between these two regions were *Z* = 0.51 ± 0.19 before the removal of EO-EPI, and *Z* = 0.19 ± 0.23 after the removal of EO-EPI (paired T(13) = 8.47, *p* < 0.001 *d* = 0.39). Additionally, the average correlation between EO-EPI and visual ROI was −0.002 ± 0.130, and sensorimotor ROI was −0.019 ± 0.120.

The above findings were obtained when the NBS method was applied at a high threshold, meaning that the edges showed a significant difference in connectivity with a threshold value exceeding *T* = 7.5. The NBS algorithm can find difference networks where the edge-level difference in connectivity strength is set at any arbitrary value. We applied the algorithm to determine if a different network would be identified at a lower threshold so that the edge difference exceeded *T* = 5.0. In this case, NBS identified difference network consisting of 133 connections (see Figure 3). For descriptive purposes, we matched each of the ROIs identified as being within the difference network to larger-scale networks in the Schaefer atlas which includes 7 networks: Visual, Somatomotor, Dorsal Attention, Ventral Attention, Limbic, Frontoparietal, and Default Mode. Of the 133 edges, 108 were between nodes of visual and somatomotor networks, ten were between the visual cortex and prefrontal cortex (assigned to DMN), three were with the temporal cortex (DMN), one was with the prefrontal cortex (control network), and two were with the operculum (salience/attention network). The Somatomotor area, on the other hand, had 2 connections with the temporal cortex (DMN), one with the operculum (salience/attention), and one with orbito-frontal cortex (limbic network). Finally, the DMN had four weaker intra-network connections and a single inter-network connection with the posterior dorsal attention network (in the prefrontal cortex).

**Figure 3.**
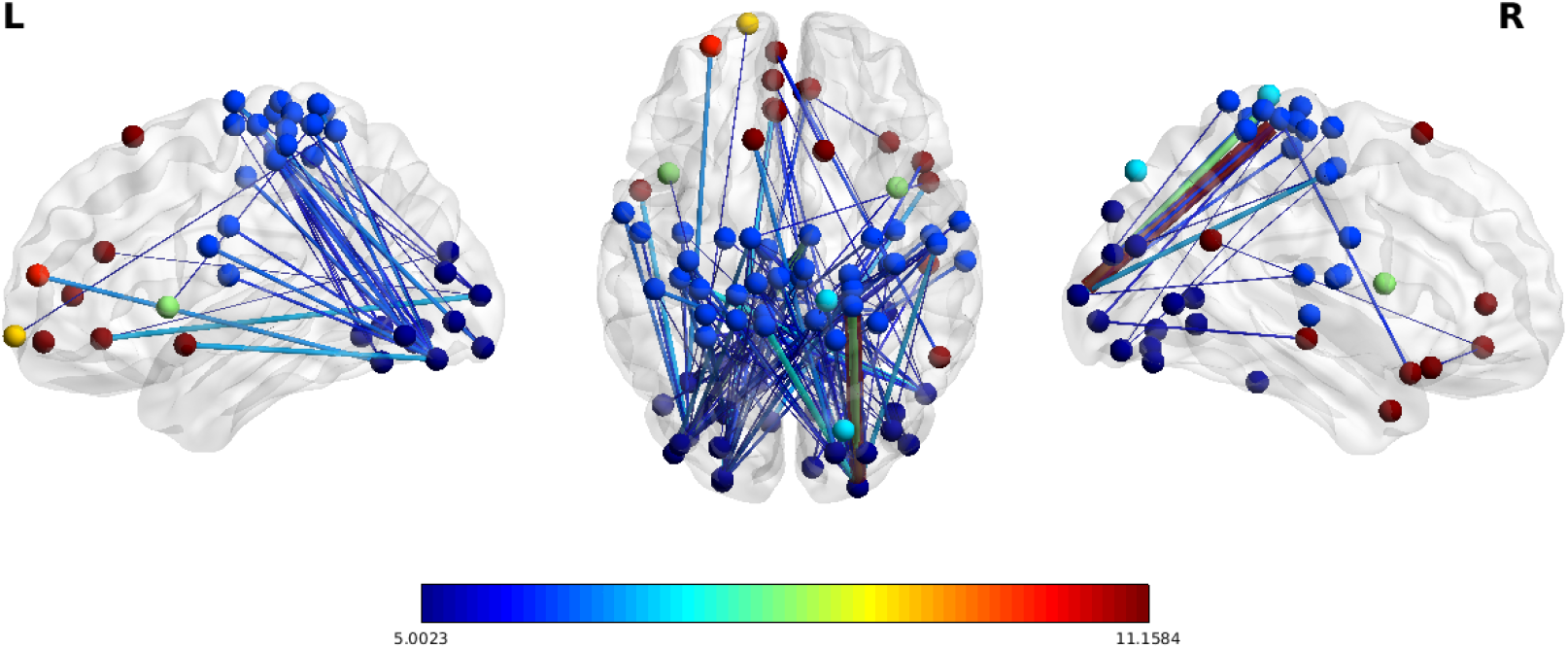
Brain areas demonstrating weaker connectivity after removal of eye-movement-related variance. A difference network was identified when edge-level differences were thresholded at *T* = 5.0. Each of the edges reflects two brain areas more weakly connected after removing variance associated with eye-movement, as operationalized via EO-EPI time series. Of the 133 edges identified, 108 were between visual and somatomotor areas. Dots represent the location of the ROIs and are color-coded according to macro-scale resting-state networks. Edges represent the connections between ROIs that survive the threshold and are color-coded in relation to the magnitude of the effect associated with the removal of eye-movement contribution.

#### Effective connectivity

The Granger causality analysis of the three selected ROIs from *i*) VisCtx, *ii*) SMCtx and *iii*) EYE_*raw*_ revealed significant causality from EYE_*raw*_ to the VisCtx (mean p-value across participants; *p* = 0.03) and from the VisCtx to the SMCtx (mean *p* = 0.02). All other tests (SMCtx to EYE_*raw*_ (*p* = 0.13), SMCtx to VisCtx (*p* = 0.17), VisCtx to EYE_*raw*_ (*p* = 0.2), and EYE_*raw*_ to SMCtx (p = 0.13) showed p values greater than 0.05 (see Figure 7).

## Discussion

Our findings indicate that hard-wired ontogenetic processes can delineate functional networks that are resistant to changes, even if they do not serve an obvious function. An ocular-related connection between the visual and sensorimotor cortices is functional in sighted, but appears to reflect a residual organization with no apparent function in the CB.

Here, we studied whether spontaneous eye movements in CB individuals are coordinated with brain activity. The congenitally blind lack a vestibulo-ocular reflex, cannot initiate voluntary saccades and may display non-coordinated eye movements. For these reasons, it was not at all given that coordinated patterns of brain activity would accompany CB’s eye movements. Furthermore, even if eye movements were coordinated with brain activity at the single participant level, there was no guarantee these would show consistency at the group level.

The EO-EPI signal was used as a proxy for eye movements (following, e.g. Beauchamp, 2003; Frey et al., 2021; Koba et al., 2021; Schöpf et al., 2014). Consistent with prior reports (e.g., Kompf & Piper, 1987; Leigh & Zee, 1980), differences between oculomotor dynamics in CB and sighted were found, with CB showing greater variance in the time series overall (Figure 1B) and increased power in both the relatively higher and lower frequency ranges. A few time series exhibited highly regular oscillatory dynamics (Figure 1A), and a few CB participants showed uncorrelated eye movements.

At the group level, we identified brain activity that correlated with the EO-EPI data, for both the raw and convolved versions. As indicated by the Dice coefficient, the distribution of this activity overlapped more strongly with that of sighted when using the raw regressor. We note that given the dynamics of CB’s EO-EPI time series, the raw time series could correlate with brain activity if peaks in the eye movement follow movement onsets by a 2-6 seconds delay, which is consistent with the slow oscillatory dynamics evident in Figure 1A. EO-EPI signal was correlated with brain activity in regions including the cerebellum, visual cortex, sensorimotor areas bilaterally, supplementary-motor area, basal ganglia and thalamus.

The cerebellar finding is particularly interesting, as this region is considered crucial to the control of eye movement. Multiple cerebellar subregions are involved in controlling different aspects of eye movements: Crus I and Crus II for control of smooth pursuit eye movements, lateral cerebellum for control of saccadic eye movements, and posterior lobe for integration of sensory information from the visual and vestibular systems to coordinate eye movements (reviewed in Kheradmand & Zee, 2011; Shemesh & Zee, 2019). For CB we find extensive positive correlations in the posterior cerebellum, potentially suggesting an intact integration of ocular and vestibular information at the cerebellar level, even if lacking a vestibu-ocular reflex.

The fact that EO-EPI is correlated with overlapping brain areas in CB and sighted (quantified from Koba et al., 2021) indicates that eye movements may play a role in mediating resting-state brain connectivity, regardless of visual input and visual experience. It is known that visually-driven stimulation of the visual system in the first two years of life is required for the normal maturation of the visual pathway and its integration with the rest of the brain (reviewed in Fine and Park, 2018 and Voss, 2019). Congenital visual deprivation and sight-recovery studies report that in the absence of early post-natal visual experience, the visual system shows irreversible functional and structural changes such as increased cortical thickness and cross-modal responses in the primary visual cortex (Collignon et al., 2015; Guerreiro et al., 2015; Hölig et al., 2022; Saenz et al., 2008). The presence of eye-movement-mediated cortical activity in blind individuals, therefore, instead suggests the existence of a purely physiological constraint on connectivity, which may reflect an initial, non-pruned state rather than one shaped by perceptual input. This is consistent with the findings of Schöpf et al. (2014), who observed correspondence between EO-EPI and resting state data in pre-natal infants.

We examined the impact of removing the EO-EPI variance on the topography and the causal structure of brain connectivity. A ‘difference network’ analysis identified a set of interconnected regions in CB whose connections were all reduced after EO-EPI removal. As shown in Figures 3 and 7, these included visual and sensorimotor areas. Using Granger analysis, we found that eye-movement data tended to precede activity in the visual area, which in turn preceded activity in the sensorimotor area.

From one perspective, the consistency of this functional connectivity pattern provides further support for existing theoretical models that suggest an innate presence of a large-scale topographic organization that maintains the functional development and the neural makeup of the deprived cortical areas (Heimler & Amedi, 2020; Ricciardi et al., 2014; Setti et al., 2023). This modality-invariant brain organization maintains, as seen in this instance, coordinated activity between brain areas whose functional role in congenitally blind people might appear ambiguous, or even ‘nonfunctional’ when viewed in the limited context of oculomotor activity. Nonetheless, this preserved interaction between visual and sensorimotor systems could conversely provide some suggestions on the functional ‘fate’ of the primary visual areas that, in congenitally blind people, respond to non-visual modalities across various tasks following organizational principles similar to those implemented for vision in sighted individuals (reviewed in Castaldi et al., 2020; Voss, 2019). Indeed, we cannot exclude that the deprived visual cortex may still have the capacity to analyze stimulus characteristics that are conveyed via non-visual inputs, and to provide task-related outputs to other sensory structures (Bridge & Watkins, 2019; Heimler & Amedi, 2020; Ricciardi et al., 2014).

## Conclusions

Our findings are the first to show that eye movement activity in CB is systematically linked to patterns of brain activity in both cortical and cerebellar areas, and furthermore, as for sighted, eye movement activity is associated with coordinated activity between visual and sensorimotor regions. These findings support the idea that this ocular/visual/sensorimotor circuit is under tight genetic control, and that its emergence and maintenance is experience independent.

## Authors and Author contributions statement

U.H, C.K, E.R, and O.C conceived the experiment(s), U.H, C.K., and A.C determined the analysis protocol, C.K, and A.C. conducted data analysis, O.C contributed neuroimaging data, U.H and C.K drafted the manuscript, All authors reviewed and edited the draft.

## Supporting Information

**Supplementary Table 1.** Areas whose resting-state BOLD activity correlated with EYE_*raw*_ as determined by the AAL2, Desikan-Killiany, and Harvard-Oxford atlases. Peak value represents 1 – *alpha*, so that values closer to 1 indicate higher statistical significance. Negative values indicate the region showed a negative relationship between RS activity and EYE_*raw*_.

| cluster_id | peak_x | peak_y | peak_z | peak_value | volume_mm | aal | desikan_killiany | harvard_oxford |
| --- | --- | --- | --- | --- | --- | --- | --- | --- |
| 1 | 65.5 | -36.5 | 41.5 | -0.972 | 200196 | SupraMarginal_R | Unknown | 27.0% Right_Supramarginal_Gyrus_posterior_division |
| 2 | -27.5 | -84.5 | -42.5 | 0.9652 | 39132 | Cerebellum_Crus2_L | Left-Cerebellum-Cortex | 0% no_label |
| 3 | -48.5 | -69.5 | 13.5 | 0.9626 | 21636 | Temporal_Mid_L | Left-Cerebral-White-Matter | 45.0% Left_Lateral_Occipital_Cortex |
| 4 | 26.5 | 32.5 | 49.5 | 0.964 | 7992 | Frontal_Sup_2_R | Unknown | 31.0% Right_Superior_Frontal_Gyrus; 29.0% Right_Middle_Frontal_Gyrus |
| 5 | 5.5 | 26.5 | -6.5 | 0.9596 | 6660 | Cingulate_Ant_R | Right-Cerebral-White-Matter | 19.0% Right_Subcallosal_Cortex |
| 6 | -24.5 | 17.5 | 41.5 | 0.953 | 6516 | Frontal_Mid_2_L | ctx-lh-superiorfrontal | 33.0% Left_Superior_Frontal_Gyrus; 13.0% Left_Middle_Frontal_Gyrus |
| 7 | 50.5 | 17.5 | 21.5 | 0.9638 | 5364 | Frontal_Inf_Tri_R | ctx-rh-parsopercularis | 52.0% Right_Inferior_Frontal_Gyrus_pars_opercularis |
| 8 | -36.5 | 20.5 | -18.5 | 0.9616 | 4212 | OFCpost_L | Left-Cerebral-White-Matter | 68.0% Left_Frontal_Orbital_Cortex |
| 9 | 53.5 | -69.5 | 29.5 | 0.9596 | 3456 | Angular_R | ctx-rh-inferiorparietal | 53.0% Right_Lateral_Occipital_Cortex_superior_division |
| 10 | 2.5 | -33.5 | 41.5 | 0.96 | 2268 | Cingulate_Mid_R | Unknown | 87.0% Right_Cingulate_Gyrus_posterior_division |
| 11 | -45.5 | 53.5 | -10.5 | 0.957 | 1944 | no_label | Unknown | 31.0% Left_Frontal_Pole |
| 12 | -12.5 | 50.5 | -10.5 | 0.9582 | 1152 | no_label | Left-Cerebral-White-Matter | 17.0% Left_Frontal_Medial_Cortex |
| 13 | -6.5 | -60.5 | 45.5 | 0.9544 | 864 | Precuneus_L | ctx-lh-precuneus | 72.0% Left_Precuneous_Cortex |

**Supplementary Figure 1.**
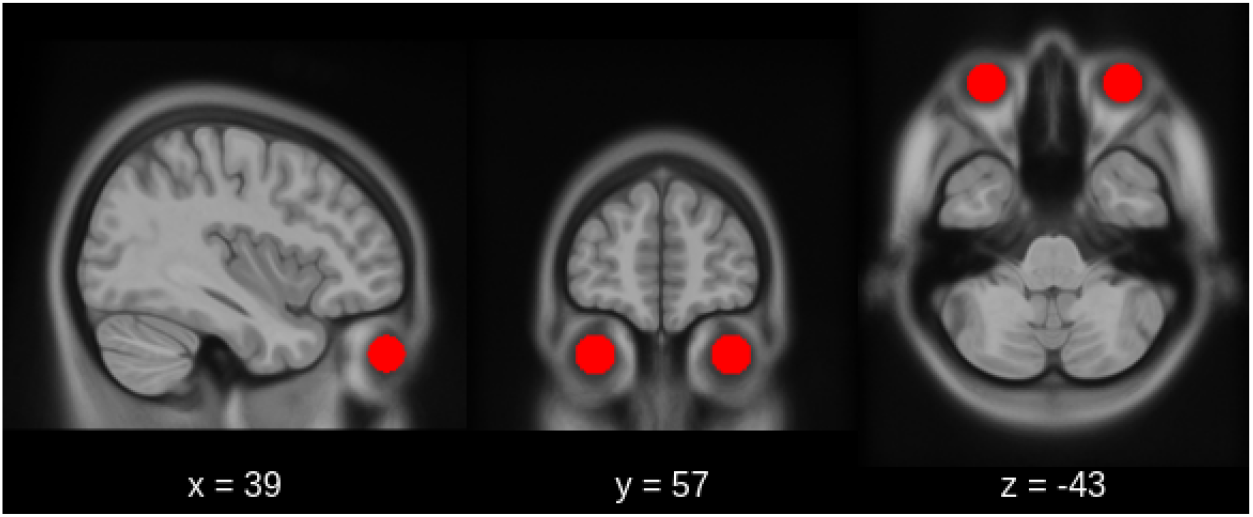
Eo-epi mask (shown in red) as drawn on the common template.

**Supplementary Figure 2.**
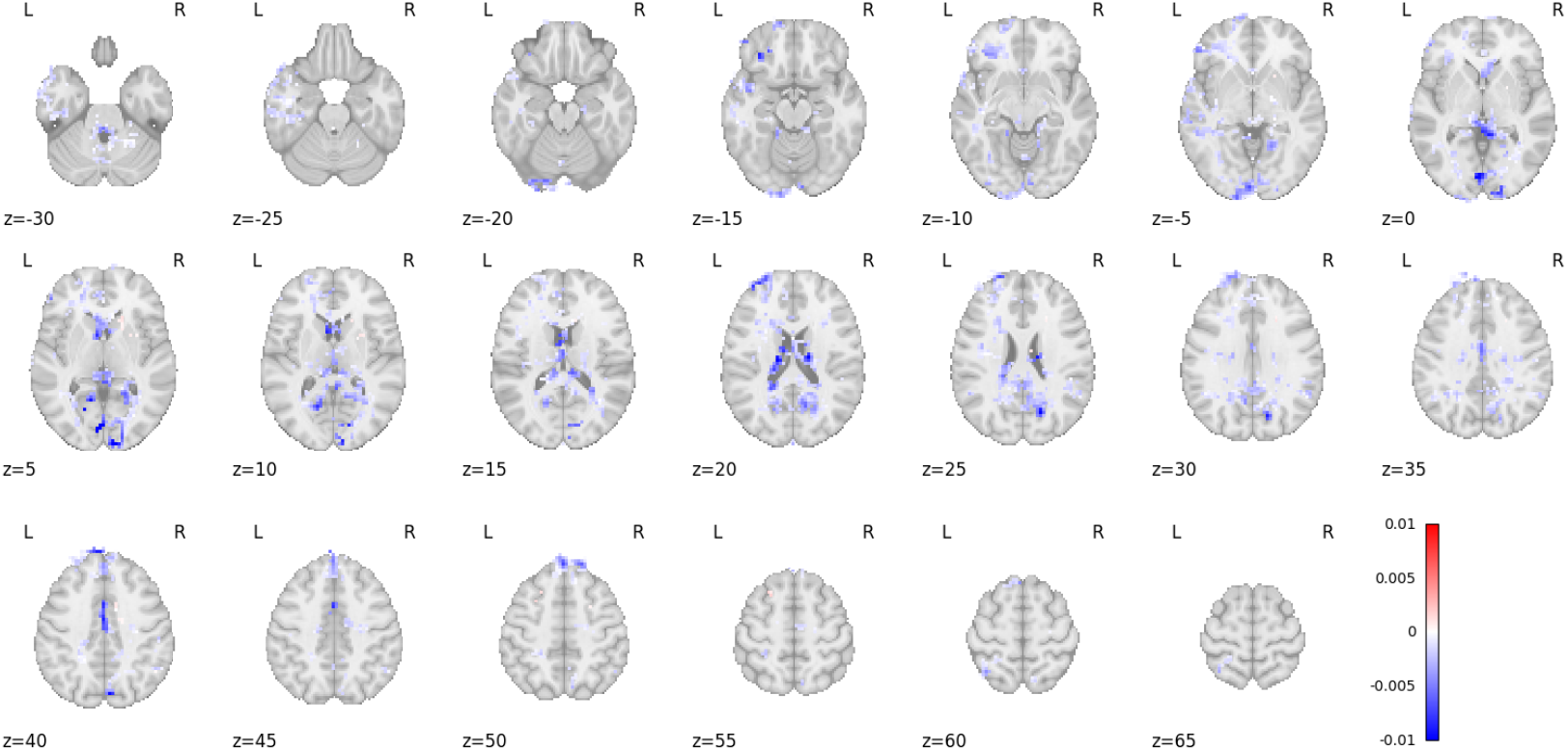
Correlates of EYE_conv_, p<0.05, corrected with TFCE. Warm colors represent a positive relationship between RS and EYE_conv_; while cold colors represent a negative one.

**Supplementary Table 2.** Areas whose resting-state BOLD activity correlated with EYE_conv_ as determined by the AAL2, Desikan-Killiany, and Harvard-Oxford atlases. Peak value represents 1 – *alpha*, so that values closer to 1 indicate higher statistical significance. Negative values indicate the the region showed negative relationship between RS activity and EYE_*conv*_.

| cluster_id | peak_x | peak_y | peak_z | peak_value | volume_mm | aal | desikan_killiany | harvard_oxford |
| --- | --- | --- | --- | --- | --- | --- | --- | --- |
| 1 | -0.5 | -9.5 | -2.5 | -0.9584 | 101664 | no_label | 3rd-Ventricle | 24.0% Left_Thalamus |
| 2 | -42.5 | -15.5 | -38.5 | -0.9552 | 8172 | no_label | Unknown | 28.0% Left_Inferior_Temporal_Gyrus; 20.0% Left_Temporal_Fusiform_Cortex |
| 3 | -6.5 | -36.5 | -34.5 | -0.9586 | 4752 | no_label | Brain-Stem | 99.0% Brain-Stem |
| 4 | 2.5 | -3.5 | 33.5 | -0.953 | 3132 | Cingulate_Mid_R | Unknown | 90.0% Right_Cingulate_Gyrus_anterior_division |
| 5 | 53.5 | -54.5 | 29.5 | -0.9524 | 2304 | Angular_R | Right-Cerebral-White-Matter | 73.0% Right_Angular_Gyrus; 10.0% Right_Lateral_Occipital_Cortex |
| 6 | -39.5 | -12.5 | 25.5 | -0.9512 | 2196 | no_label | Left-Cerebral-White-Matter | 0% no_label |
| 7 | -33.5 | -39.5 | -6.5 | -0.9512 | 1872 | no_label | Left-choroid-plexus | 16.0% Left_Lateral_Ventrical; 13.0% Left_Parahippocampal_Gyrus |
| 8 | -57.5 | -24.5 | -2.5 | -0.955 | 1836 | Temporal_Mid_L | Unknown | 38.0% Left_Superior_Temporal_Gyrus; 12.0% Left_Middle_Temporal_Gyrus |
| 9 | 17.5 | -54.5 | -42.5 | -0.9534 | 1260 | Cerebellum_8_R | Right-Cerebellum-White-Matter | 0% no_label |

**Supplementary Figure 3.**
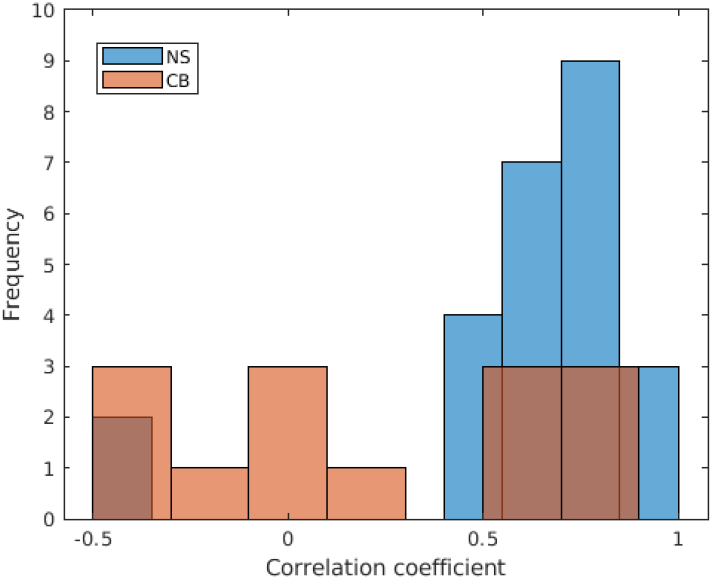
Histogram of the correlation values between right and left EO-EPI of CB and normally Sighted (NS). Correlations for Sighted are more consistent and show higher positive values.

**Supplementary Figure 4.**
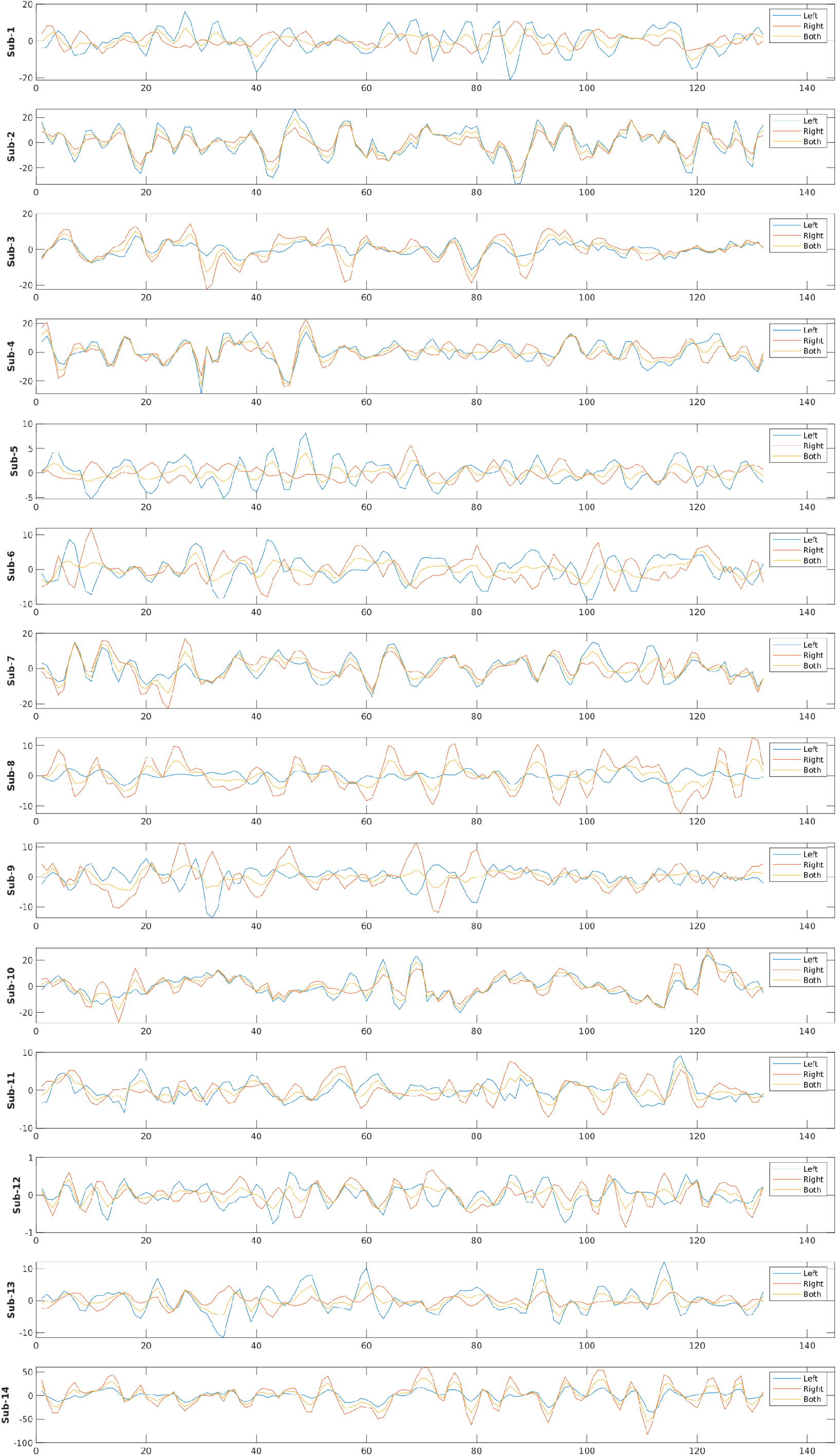
EO-EPI from right and left eyes, and their average from all subjects.

**Supplementary Figure 5.**
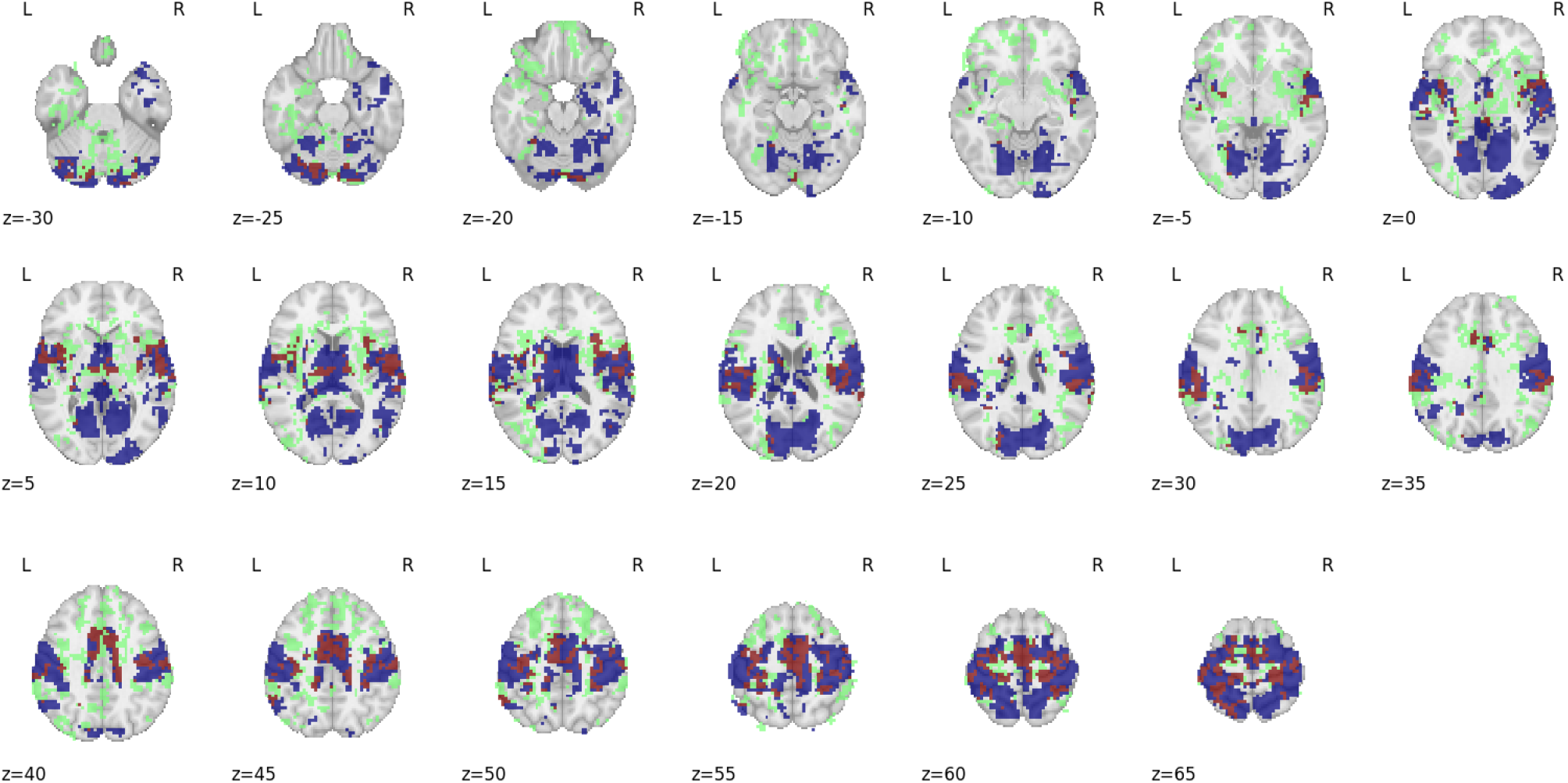
Overlap between brain areas correlating with eye movements in Congenitally Blind in the current study and normally Sighted. Data from Sighted were collected by Koba et al. 2021. Red areas indicate areas correlated with eye movements for both groups. Blue areas were identified for Sighted alone (N=83), and green areas for Blind alone (N=14). As evident in the figure, both groups show bilateral eye-movement related activity in sensorimotor cortex and visual cortices. Congenital Blind participants appear to show greater involvement of the frontal cortex

**Supplementary Figure 6.**
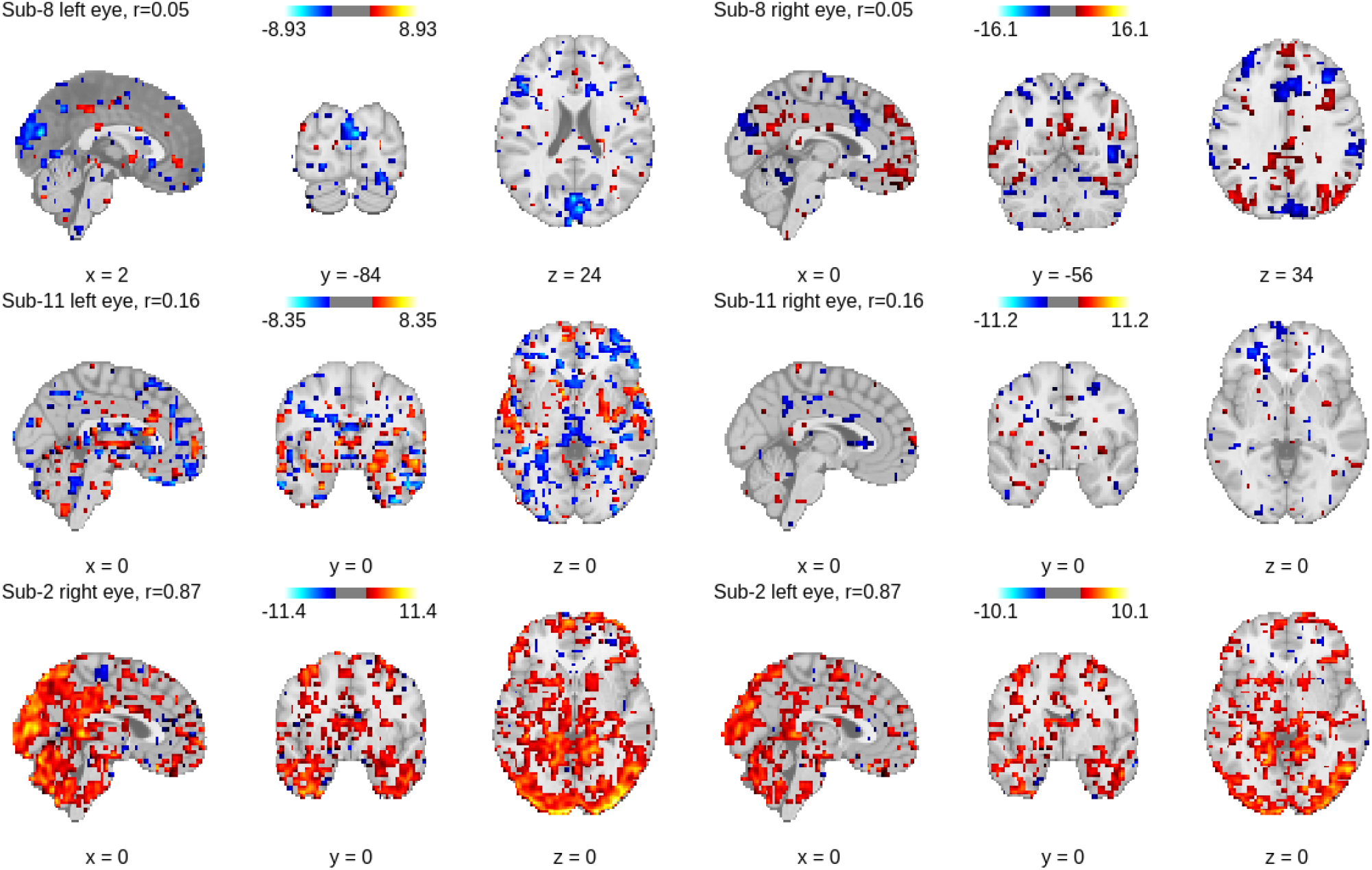
Relationship between right or left EO-EPI and RS for sample participants. First two rows present participants whose eyes demonstrate low EO-EPI correlation, and the bottom row demonstrates a participant with high correlation. Images are thresholded at an arbitrary threshold of t =2.6. It can be seen that right and left eyes pick up similar regions when the correlation between eyes is high, and when the correlation is low, they pick up different regions.

**Supplementary Figure 7.**
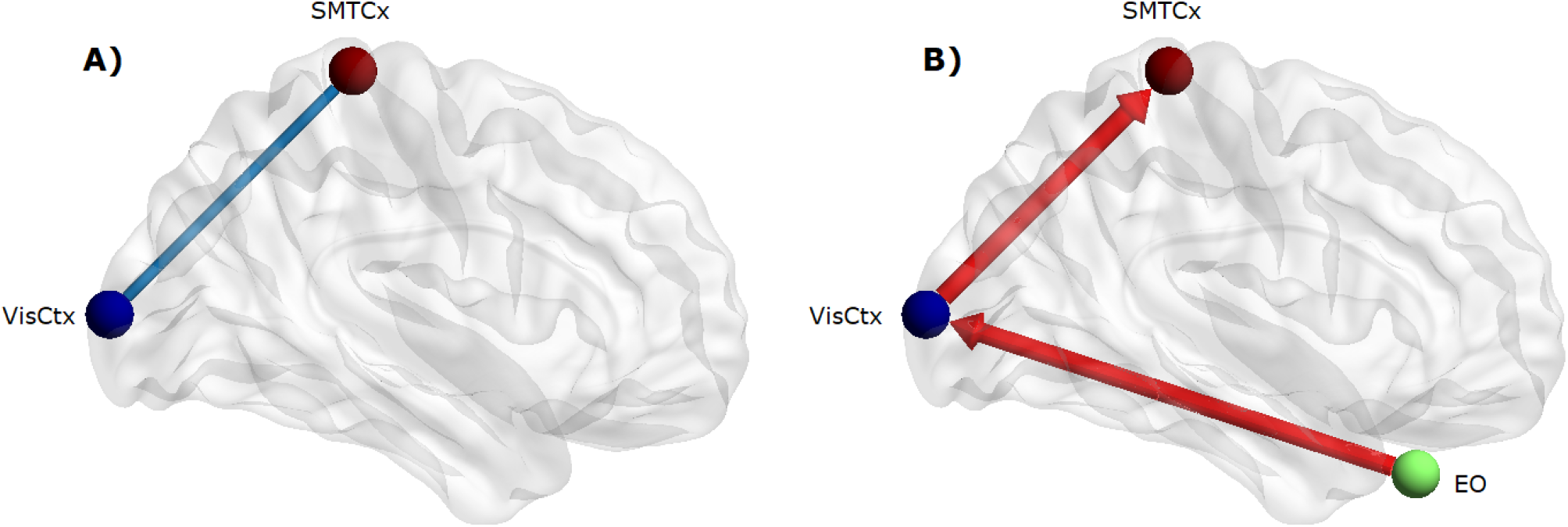
**A)** Weaker functional connectivity found in the RS networks cleaned from EYE_*raw*_, compared to regular RS networks. The nodes represent the visual cortex (roi name: RH_Vis_21, blue) and the sensorimotor cortex (roi name: RH_Sommot_35, red). **B)** Granger causality tests showed significant causal effects of EYE_*raw*_ on visual cortex (*p* =0.03) as well as of visual cortex on the sensorimotor area (*p* =0.02). The remaining relationships among those three regions were not significant (*p* values lower than 0.05).

## Notes

### Competing Interest Statement

The authors have declared no competing interest.

### Summary of Updates

Author order updated

